# APART-QSM: an improved sub-voxel quantitative susceptibility mapping for susceptibility source separation using an iterative data fitting method

**DOI:** 10.1101/2023.04.02.535256

**Authors:** Zhenghao Li, Ruimin Feng, Qiangqiang Liu, Jie Feng, Guoyan Lao, Ming Zhang, Jun Li, Yuyao Zhang, Hongjiang Wei

**Affiliations:** School of Biomedical Engineering, Shanghai Jiao Tong University, Shanghai, China; Department of Neurosurgery, Clinical Neuroscience Center Comprehensive Epilepsy Unit, Ruijin Hospital, Shanghai Jiao Tong University School of Medicine, Shanghai, China; School of Information Science and Technology, ShanghaiTech University, Shanghai, China

**Keywords:** Susceptibility source separation, QSM – quantitative susceptibility mapping, Paramagnetic, Diamagnetic

## Abstract

The brain tissue phase contrast in MRI sequences reflects the spatial distributions of multiple substances, such as iron, myelin, calcium, and proteins. These substances with paramagnetic and diamagnetic susceptibilities often colocalize in one voxel in brain regions. Both opposing susceptibilities play vital roles in brain development and neurodegenerative diseases. Conventional QSM methods only provide voxel-averaged susceptibility value and cannot disentangle intravoxel susceptibilities with opposite signs. Advanced susceptibility imaging methods have been recently developed to distinguish the contributions of opposing susceptibility sources for QSM. The basic concept of separating paramagnetic and diamagnetic susceptibility proportions is to include the relaxation rate 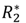 with 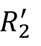 in QSM. The magnitude decay kernel, describing the proportionality coefficient between 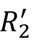 and susceptibility, is an essential reconstruction coefficient for QSM separation methods. In this study, we proposed a more comprehensive complex signal model that describes the relationship between 3D GRE signal and the contributions of paramagnetic and diamagnetic susceptibility to the frequency shift and 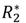 relaxation. The algorithm is implemented as a constrained minimization problem in which the voxel-wise magnitude decay kernel and sub-voxel susceptibilities are determined alternately in each iteration until convergence. The calculated voxel-wise magnitude decay kernel could realistically model the relationship between the 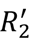 relaxation and the volume susceptibility. Thus, the proposed method effectively prevents the errors of the magnitude decay kernel from propagating to the final susceptibility separation reconstruction. Phantom studies, *ex vivo* macaque brain experiments, and *in vivo* human brain imaging studies were conducted to evaluate the ability of the proposed method to distinguish paramagnetic and diamagnetic susceptibility sources. The results demonstrate that the proposed method provides state-of-the-art performances for quantifying brain iron and myelin compared to previous QSM separation methods. Our results show that the proposed method has the potential to simultaneously quantify whole brain iron and myelin during brain development and aging.

The proposed model was also deployed with multiple-orientation complex GRE data input measurements, resulting in high-quality QSM separation maps with more faithful tissue delineation between brain structures compared to those reconstructed by single-orientation QSM separation methods.

## 1. Introduction

Quantitative susceptibility mapping (QSM) is a technique based on MRI phase signal that quantifies the spatial distribution of magnetic susceptibility within a measured tissue (de Rochefort et al., 2008; Haacke et al., 2015; Liu et al., 2015; Shmueli et al., 2009; Wang and Liu, 2015). The susceptibility contributors include biometals and molecules, e.g., iron, myelin, calcium, and lipids (Bilgic et al., 2012; Deistung et al., 2017; He et al., 2015; Li et al., 2011; Reichenbach et al., 2015; Schweser et al., 2012; Sun et al., 2018; Wei et al., 2016). The temporal changes of iron content, myelin, and proteins are involved in normal brain development and a variety of neurodegenerative diseases, e.g., iron overload in Parkinson’s disease (PD) (Acosta-Cabronero et al., 2017; Du et al., 2016; Guan et al., 2019; He et al., 2015; Uchida et al., 2019), demyelination in Multiple Sclerosis (MS) (Kaunzner et al., 2019; Li et al., 2016; Schweser et al., 2018) and amyloid β-protein accumulation in Alzheimer’s disease (AD) (Acosta-Cabronero et al., 2013; Ayton et al., 2017; Cogswell et al., 2021; Gong et al., 2019; Spotorno et al., 2020). QSM has shown great potential for quantifying the magnetic susceptibility of these substances in the past decade.

In the central nervous system (CNS), multiple neurophysiological processes with various substances might co-occur in local brain regions because of the limited spatial resolution in current MRI. For instance, iron deposition is involved in many important processes such as myelin synthesis and maintenance, myelin production and metabolism of neurotransmitters. The tight association of iron deposition with axonal myelination has been reported in brain development (Moller et al., 2019; Ward et al., 2014). In various neurodegenerative diseases, abnormal iron homeostasis can cause the modification of lipids and proteins. For example, metal-binding to amyloid β is likely to be involved in the pathology of AD (Becerril-Ortega et al., 2014; Wärmländer et al., 2019). Iron deposition has been widely suspected as the underlying cause of AD, aggregating with amyloid β. However, traditional QSM methods cannot differentially quantify molecular sources with opposing susceptibility sources, such as paramagnetic iron and diamagnetic amyloid β, within the same voxel. The paramagnetic and diamagnetic susceptibility effects may cancel out partly or in whole, resulting in an underestimated susceptibility value and inaccurate quantification results. Therefore, it is highly desirable to disentangle the underlying susceptibility proportions and to provide a more specific quantification of tissue magnetic properties.

Previous studies have attempted to separate the contributions of paramagnetic and diamagnetic susceptibility sources for QSM. The first relevant study was made by Schweser et al., who tried to separate iron and myelin concentrations by assuming both 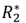 and susceptibility values depend linearly on iron and myelin concentration (Schweser et al., 2011). The coefficients were pre-determined by offline calibration experiments on 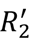, susceptibility and magnetization transfer saturation images. It is noteworthy that the coefficients were assumed to be equal for all types of brain tissue. Stüber et al. proposed to quantify the iron and myelin concentrations by linear regression analysis between the 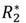 and magnetic susceptibility versus iron and myelin concentrations inferred by MRI signal and proton-induced X-ray emission in post-mortem brain tissues (Stüber et al., 2014). Recently, χ -separation (Lee et al., 2017; Shin et al., 2021) allows the separation of opposing susceptibility sources by assuming that frequency shift linearly depends on the composition of susceptibilities and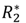 is linearly affected by the absolute value of susceptibilities. 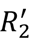 is a fundamental precondition for the feasibility of the separation of opposing susceptibility sources. The magnitude decay kernel, representing the proportionality constant between 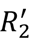 and the absolute susceptibility, is an essential reconstruction coefficient for χ -separation. The decay kernel was pre-calculated as the slope of the linear regression of the mean 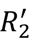 with respect to the susceptibility values from a few deep gray matter (DGM) regions and was then applied to the whole brain as a spatially invariant coefficient (Shin et al., 2021; Zhang et al., 2022). Similarly, Emmerich et al. determined the value of the magnitude decay kernel by a phantom study and applied this value to the experimental human brain data for the separation of susceptibility sources (Emmerich et al., 2021). However, the proportion of various susceptibility sources may vary across brain DGMs and contribute to the field inhomogeneity and 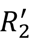 differently for different voxels. Thus, the assumption of the spatially invariant coefficient (i.e., the magnitude decay kernel) used to differentiate the paramagnetic susceptibility from the diamagnetic susceptibility sources could be erroneous. Chen et al. proposed to decompose volume susceptibility into sub-voxel paramagnetic and diamagnetic components based on a three-pool complex signal model, named DECOMPOSE-QSM (Chen et al., 2021). DECOMPOSE-QSM relies on the pre-calculated QSM maps at different echo times to synthesize the signal instead of using the raw complex GRE signal or tissue phase maps as input. The linear coefficient between 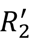 and the single susceptibility source was also assumed to be spatial invariant with the value of 323.5 Hz/ppm at 3.0 T MRI under the static dephasing regime assumption. However, the assumption of the relaxation following the static dephasing regime theory might be void, especially in brain white matter due to highly anisotropic axon and myelin (Duyn and Schenck, 2017).

In this study, inspired by χ-separation, we introduced a more comprehensive model to separate the paramagnetic and diamagnetic susceptibilities within the voxel by combining 3D gradient recalled echo (GRE) data with T2 mapping. The QSM dipole inversion was inherently integrated into the proposed model. The opposing susceptibility values within the sub-voxel can be directly quantified by fitting the acquired data to the proposed complex-valued model. Additionally, starting with an initial estimation of the magnitude decay kernel and susceptibilities, the method alternately updated the reconstructed susceptibilities and the magnitude decay kernel in each iteration and repeated until convergence for each voxel. The proposed method was termed iterAtive magnetic suscePtibility sources sepARaTion or APART-QSM. APART-QSM was validated using a phantom experiment and *ex vivo* macaque brain experiments with myelin staining as the ground truth. Finally, our method was applied to 32 healthy subjects with age from 4 to 39 years old to explore the temporal trajectories of paramagnetic and diamagnetic susceptibilities in the human brain DGMs. The results demonstrated the superiority of our proposed method in terms of susceptibility quantification accuracy, indicating its further applications for the investigation of myelin-and iron-related diseases. Additionally, the proposed APART-QSM was deployed with multiple-orientation complex GRE data input measurements, resulting in high-quality QSM separation maps with improved tissue delineation between brain structures compared to those reconstructed by single-orientation QSM separation methods. Our proposed method has three major differences from previous studies, i.e., χ-separation and DECOMPOSE-QSM. (1) We proposed a more comprehensive complex signal model that describes the relationship between 3D GRE signal and the contributions of paramagnetic and diamagnetic susceptibility to the frequency shift and 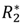 relaxation. (2) We alternately estimated the magnitude decay kernel and the reconstructed susceptibilities in an iterative fitting fashion for each voxel, providing state-of-the-art reconstruction results of susceptibility separation. (3) The proposed method can handle an arbitrary number of complex GRE data input measurements to provide high-quality QSM separation maps with more faithful tissue delineation of the small brain sub-regions.

## 2. Theory

### 2.1 Biophysical model of susceptibility source separation

The materials in a static magnetic field *B*_0_ can be excited by a specific radio frequency (RF) pulse, resulting in an *R_2_* transverse relaxation. Due to the field inhomogeneity caused by external susceptibility sources, an extra transverse relaxation 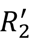 in position *r* will be produced under the static dephasing regime (Haacke et al., 1989; Yablonskiy and Haacke, 1994):

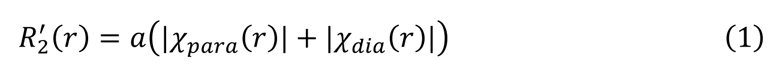

where 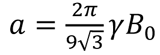 is the magnitude decay kernel which is a bulk proportionality constant between 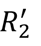 and absolute values of susceptibility sources. *γ* is the

gyromagnetic ratio. In brain tissues, χ_*para*_ and χ_*dia*_ denote the paramagnetic and diamagnetic susceptibility sources which are mostly dominant by iron and myelin, respectively. Eq. (1) reveals that 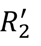 can be mathematically divided into 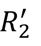_*para*_ and 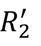_*dia*_, which are linearly dependent on the paramagnetic and diamagnetic susceptibility sources within a voxel.

The magnetic susceptibility χ induces extra field inhomogeneities, causing a detectable frequency shift Δ*f* by MRI (Deistung et al., 2017). χ can also be expressed as the signed sum of two opposing susceptibility sources. Thus, the frequency shift caused by magnetic susceptibilities can be written as:

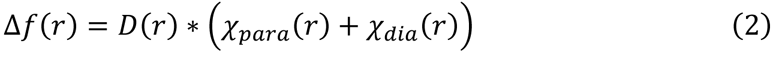

where *D* is the magnetic dipole kernel and ∗ denotes the spatial convolution.

The susceptibility separation can be modeled by complex MR signals by combining Eqs. (1) and (2), as proposed by Shin et al., called χ-separation (Shin et al., 2021):

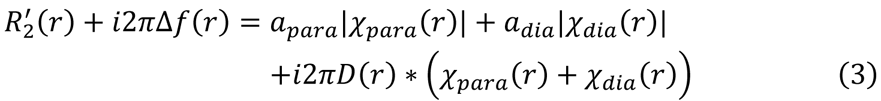

where *a_para_* and *a_dia_* were considered spatial invariances in χ-separation, and they were assumed to be equal, determined by the ratio between 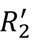 and susceptibility in a few iron-rich DGM regions.

### 2.2 The iterative complex-valued model for susceptibility separation

The raw complex GRE signal *S* can be expressed by the susceptibility sources and modeled as (details are in Appendix):

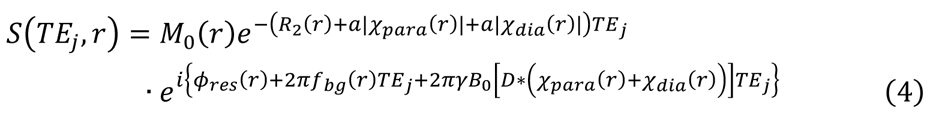

where *M*_0_ is the extrapolated magnitude signal at TE = 0 ms. ɸ_*res*_ represents the time-independent residual phase. 2Π*f*_*bg*_TE_*j*_ denotes the background phase. *j* denotes the index of echo times.

To solve the complex model, the GRE signals are separated as magnitude and phase terms and the proposed APART-QSM is eventually formulated as a two-step optimization. *M*_0_ and the relaxation rate 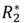 can be simultaneously estimated following an exponential model (Wang et al., 2020):

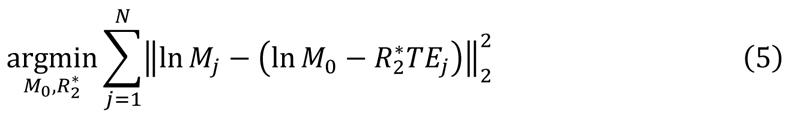

where *N* is the total number of echo times, ‖⋅‖_2_ represents *L*2 norm, and *M*_*j*_ denotes the *j*^th^ magnitude signal. Then combining the *R_2_* map, which can be obtained by an additional T2 mapping scan, χ_*para*_ and χ_*dia*_ as well as ɸ_*res*_ can be solved as follows:

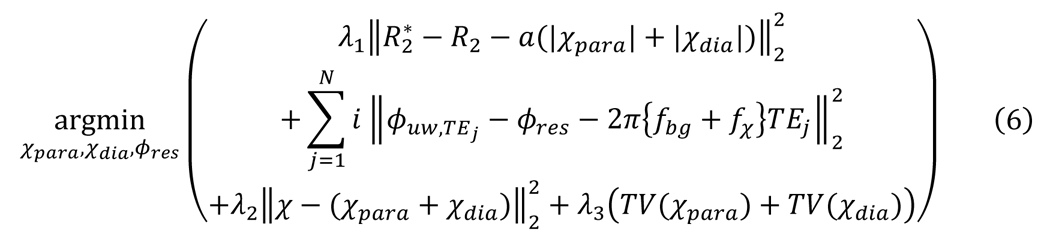

where *f*_χ_ = *γB*_0_[*D* ∗ (χ_*para*_ + χ_*dia*_)], *ϕ_uw,TEj_* is the unwrapped phase image, and χ is a pre-reconstructed QSM map providing an approximate susceptibility range constraint for the sum of the two susceptibility components. *TV(.)* is a 3D spatial total variation operator. The regularization parameters λ_1_ and λ_2_ are applied for normalizing range consistent with *ϕ*_uw,TE_*j* and *ʎ*_3_ is the regularization parameter of total variation controlling the denoising level.

The parameter *ɑ* was regarded as spatial invariant in previous studies. However, *in vivo* brain tissue contains complicated mixtures of the susceptibility sources with the non-neglectable water diffusion effect. Thus, the parameter *ɑ* in APART-QSM is considered as a voxel-specific parameter relating 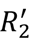 and absolute susceptibility. To address this problem, we resort to an iterative algorithm to implement the proposed APART-QSM method to fit a voxel-specific *ɑ*-map, and thus prevent the errors of parameter *ɑ* from propagating to the final susceptibility separation reconstruction.

To reconstruct the susceptibility images, we first obtain the initial estimates of the opposing susceptibilities and time-independent residual phase using Eq. (6) with an initial theoretical *ɑ* -map. These estimations may result in inaccurate susceptibility separation values due to the variable susceptibility source concentrations in different voxels. Therefore, the parameter *ɑ* needs to be updated to *ɑ*^′^ voxel-by-voxel:

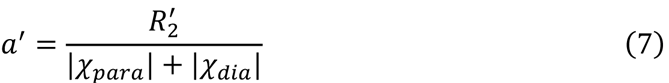

The obtained parameter *ɑ*^′^ is then plugged back into Eq. (6) to recalculate χ_*para*_ and χ_*dia*_, then obtaining *ɑ*^′^ following Eq. (7) iteratively. For the *k*^th^ iteration, the relative change of the residual *ε* in two consecutive iterations is:

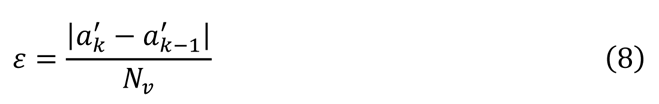

where *N*_*v*_ is the voxel number. For each iteration, Eq. (6) was implemented using a conjugate gradient-based LSQR solver. We alternate between updating the susceptibility values (χ_*para*_, χ_*dia*_) and residual phase (ɸ_*res*_) and updating the parameter *ɑ* based on Eqs. (6) and (7). The algorithm was terminated when *ε* fell below the tolerance of 0.3 Hz/ppm or the iterations reached 100. The corresponding image reconstructed at the final iteration gives the desired images.

### 2.3 Susceptibility separation from the multiple-orientation data for segmentation

The dipole inversion process was integrated into the proposed APART-QSM, as shown in Eq. (6). However, the reconstructed susceptibility maps estimated by using the iterative algorithm still suffer from streaking artifacts due to the ill-posed nature of QSM reconstruction. This issue can be overcome by using phase images acquired at different head orientations to yield susceptibility maps free of streaking artifacts, called Calculation Of Susceptibility through Multiple Orientation Sampling (COSMOS) (Liu et al., 2009). Similarly, APART-QSM was deployed with multiple-orientation complex GRE data input measurements to provide high-quality paramagnetic and diamagnetic susceptibility maps. The new optimization problem is formulated as follows:

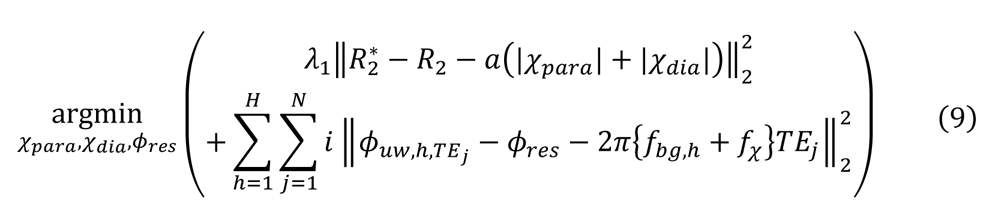

where *f*_χ_ = *γB*_0_[*D*_ℎ_ ∗ (χ_*para*_ + χ_*dia*_)], *H* is the number of orientations. *ϕ*_bg,hTEj*j*_ and 2Π*f*_bgw__*j*_ are the unwrapped phase and background phase at the ℎ^th^ head orientation, defined similarly as in Eq. (6). *D*_ℎ_ is the dipole kernel of the ℎ^th^ head orientation in the subject frame of reference. The iteration algorithm for calculating the desired susceptibilities is the same as in Section 2.2.

## 3. Materials and methods

### 3.1 Algorithm implementation

The data processing and analysis were executed on MATLAB 9.7 (R2019a MathWorks Inc., Natick, MA, USA) and deployed on a workstation with an Intel Core i7-9700 CPU and 64GB RAM. The reconstruction code and test data are available at https://github.com/AMRI-Lab/APART-QSM.

### 3.2 Agarose gel phantom fabrication and imaging

A custom cylindrical container was fabricated with a diameter of 125 mm and a height of 55 mm. A 3×3 array of small cylindrical tubes (diameter of 15 mm and height of 40 mm) was placed into the container to be filled with different concentrations of susceptibility sources. The agarose (Biogreen biotechnology) was mixed with deionized water at 1.0% w/v, and heated in a microwave oven until boiling. Then the agarose solution was poured into the container and cooled down for solidification (under 4℃). The cylindrical tubes were removed with empty cylinders. These cylindrical spaces were then filled with 1.0% w/v heated agarose solution mixed with susceptibility sources of different categories and concentrations. After these mixed agarose gels solidified, the top of the phantom was covered with the same agarose gel to avoid the signal gaps at the air-gel interface. We used iron (II, III) oxide (Fe_3_O_4_, Sinopharm Chemical Reagent Co., Ltd.) as the paramagnetic species and calcium carbonate (CaCO_3_, Sinopharm Chemical Reagent Co., Ltd.) as the diamagnetic species. The two sources were assigned with the range of susceptibility values in [-0.2, 0.2] ppm. As shown in Fig. 1A, the cylinders in the first row were filled with linearly increasing concentrations of 58.0, 116.0, and 174.0 mg/ml CaCO_3_ concentrations. The cylinders in the second row were filled with linearly increasing concentrations of 2.0, 4.0, and 6.0 μg/ml Fe_3_O_4_ concentrations. Hence, the single susceptibility sources were considered as the reference to validate the proposed model. The cylinders in the third row were filled with the gel mixed with both susceptibility sources from the first two rows.

**Figure 1.**
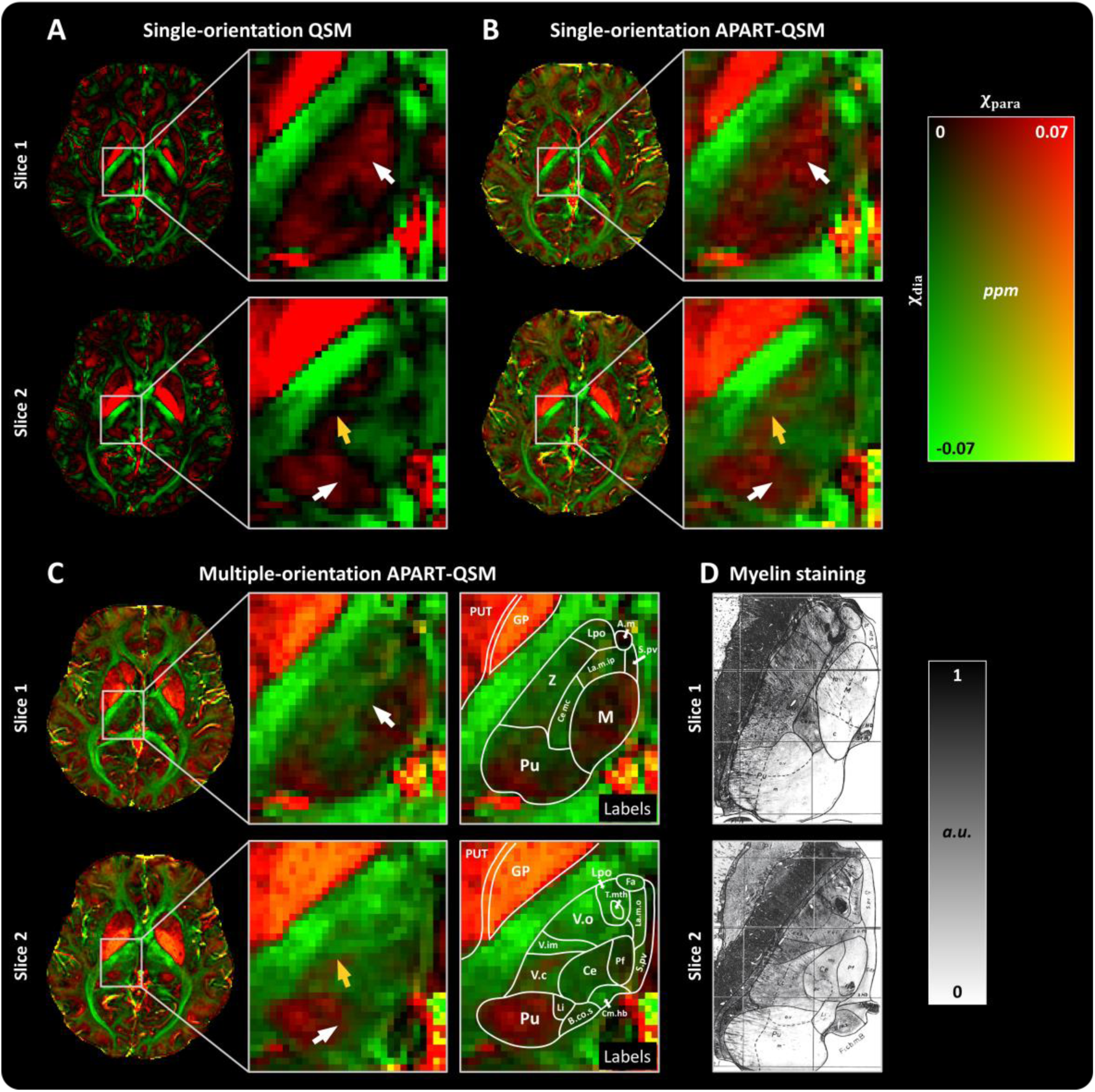
APART-QSM results of the agarose gel phantom. (A) The arrangement of phantom compositions. (B-E) Paramagnetic and diamagnetic susceptibility maps reconstructed by APART-QSM and χ-separation. (F) The voxel-specific *ɑ*-map estimated from APART-QSM. (G) Conventional QSM. The paramagnetic and diamagnetic susceptibility effects cancel out in the mixed sources cylinders in the conventional QSM. The linear regressions of single-source susceptibility versus reconstructed susceptibility from the mixed situation of each cylinder are obtained from APART-QSM (H) and χ-separation (I).

The phantom data were acquired using a 3.0 T MR scanner (uMR790, United Image Healthcare (UIH), Shanghai, China) with a 24-channel head coil. The phantom was positioned with the long axis of cylinders parallel to the main magnetic field. The 3D multi-echo GRE data were acquired with the following parameters: field of view (FOV) = 144×145×80 mm^3^; matrix size = 144×145×40; spatial resolution: 1×1×2 mm^3^; TR = 40.0 ms; TE_1_/spacing/TE_5_ = 1.9/3.2/14.7 ms; flip angle (FA) = 15°; bandwidth (BW) = 740 Hz/pixel; scan time = 5.1 min. The T2 mapping data was acquired using a 2D multi-echo spin-echo (MSE) sequence with copied FOV from 3D GRE scan: FOV = 144×145 mm^2^; matrix size = 144×145; slice number = 40; spatial resolution: 1×1×2 mm^3^; TR = 3813 ms; TE_1_/spacing/TE_5_ = 16.72/16.72/83.6 ms; BW = 160 Hz/pixel; scan time = 10.9 min.

The *R_2_* map was fitted from MSE data using a mono-exponential function in STI-Suite (https://people.eecs.berkeley.edu/-chunlei.liu/software.html). The acquired multi-echo 3D GRE phase images were first unwrapped by a Laplacian-based phase unwrapping method (Schofield and Zhu, 2003). The background phase and tissue phase were separated using the V-SHARP method (Özbay et al., 2017; Wu et al., 2012). All tissue phase images from different echoes were normalized by 2Π*γ*TE_*j*_*B*_0_ then averaged along the echo dimension. Then, STAR-QSM (Wei et al., 2015) was performed to pre-calculate the QSM map (https://people.eecs.berkeley.edu/∼chunlei.liu/software.html). The initial parameter *ɑ* was set as 323.5 Hz/ppm. Finally, the phase data, GRE magnitude data, and *R_2_* map were combined into the proposed APART-QSM to reconstruct paramagnetic and diamagnetic susceptibility maps. The regularization parameters in Eq. (6) were λ_1_ = 0.1 and λ_2_ = 10 according to the corner point in their L-curves (Fig. S3). *λ*_3_ was determined to be 1. A comparison was performed with the streaking artifact-suppressed χ-separation method, implemented on the released code. The parameters *a_para_* and *a_dia_* in χ-separation were used the same values as used in (Shin et al., 2021).

### 3.3 Ex vivo imaging and histological staining

An *ex vivo* experiment was implemented on two adult Macaca fascicularis brains for validation of APART-QSM. The brains were acquired following approved protocols from the Life Sciences Ethics Committee, Institute of Neuroscience, Chinese Academy of Sciences. The brains were used for MR scans and then stained for myelin.

The brains were immersed in 4% formalin solution for 2 months for conservation. The brain was soaked into phosphate-buffered saline (PBS) for about 1 h to recover signals before MR scanning and then transferred to an MR-compatible container, immersed in liquid fluorocarbon (Galden, PFPE, Solvay, Brussels, Belgium) for the scan. The MR data were acquired at a 9.4 T MR scanner (Bruker, BioSpec, Ettlingen, Germany). The 3D GRE scan parameters were: FOV=71.0×84.0×51.2 mm^3^; matrix size = 710×840×512; spatial resolution = 0.1×0.1×0.1 mm^3^; TR = 75 ms; TE_1_/spacing/TE_3_ = 5.7/8.5/24.7 ms; FA = 35°; BW = 81967 Hz; scan time = 6.7 h. The 3D MSE sequence was conducted with the parameters: FOV = 71.0×82.0 mm^2^; matrix size = 710×820; slice number = 512; spatial resolution = 0.1×0.1×0.12 mm^3^; TR = 140 ms; TE = 15.2, 30.4 ms; BW = 66667 Hz; scan time = 12.6 h. The DTI data were acquired by a 3D spin-echo pulse sequence: FOV = 70.7×84.0×53.2 mm^3^; matrix size = 202×240×152; spatial resolution = 0.35×0.35×0.35 mm^3^; TR = 100 ms; TE = 25.2 ms; One non-diffusion weighted image and one b-value of 3000 s/mm^2^ with 30 diffusion directions; BW = 45455 Hz; scan time = 19.9 h.

The myelin staining procedure followed the steps described in (Huitema et al., 2021), which was first dehydrated and quickly frozen by dry ice, and cut into sections 80-μm thick at the coronal position. Then the sections were stained with Luxol Fast Blue (LFB, Beijing Solarbio Science & Technology Co., Ltd) solution. The stained sections were imaged by a digital slide scanner (Axio Scan. Z1, Carl Zeiss Meditec AG, Germany).

To validate the accuracy of the estimated diamagnetic susceptibility for quantifying white matter myelin, a macaque brain parcellation (Feng et al., 2017) was used for labeling white matter bundles. For comparison, the stained myelin section was transferred to a gray map and normalized to [0, 1]. The linear regression analysis was applied to mean diamagnetic values and normalized staining values of eight white matter labels.

### 3.4 In vivo imaging and analysis

The local Human Ethics Committee approved this study and all subjects (or guardians of underage subjects) signed informed consent before scanning. 32 subjects (age = 18.3 ± 9.9 years, age range = [4, 39] years; 22 males and 10 females) were scanned on a UIH 3.0 T MR scanner (uMR890, UIH, Shanghai, China). The scan parameters of the 3D GRE were: FOV = 230×230×160 mm^3^; matrix size = 224×224×80; spatial resolution = 1.03×1.03×2 mm^3^; TR = 40 ms; TE_1_/spacing/TE_7_ = 2.4/4.3/28.2ms; FA = 15°; BW = 350 Hz/pixel; scan time = 8.7 min. The scan parameters of the 2D MSE were: FOV = 230×230 mm^2^; matrix size = 224×224; slice number = 80; spatial resolution: 1.03×1.03×2 mm^3^; TR = 3864 ms; TE_1_/spacing/TE_5_ = 16.1/16.1/80.5 ms; BW = 160 Hz/pixel; scan time = 16.1 min.

The acquired data were preprocessed and reconstructed by the proposed method to obtain the paramagnetic and diamagnetic maps, then compared to iron staining and myelin staining images of the post-mortem brains from literature (Drayer et al., 1986; Naidich et al., 2013) and website (https://brains.anatomy.msu.edu/).

Five DGM ROIs of 32 subjects were extracted by two experts manually, namely globus pallidus (GP), putamen (PUT), caudate nucleus (CN), substantia nigra (SN), and red nucleus (RN). The final labels took the intersection of ROIs from two experts. The mean paramagnetic susceptibility values in the DGMs of each subject were computed and compared with the age-related iron concentration from the post-mortem study (Aoki et al., 1989; Hallgren and Sourander, 1958) by linear regression. A multi-modality atlas was constructed using advanced normalization tools (ANTs) (Avants et al., 2010). Specifically, QSM, paramagnetic and diamagnetic susceptibility maps from all subjects were first registered to the Montreal Neurological Institute (MNI) space and then followed by a nonlinear group-wise normalization method. The diamagnetic susceptibility values of five DGMs of the atlas were compared with normalized human brain myelin sections (https://brains.anatomy.msu.edu/). Additionally, the temporal changes of DGM iron deposition and myelination were also quantified to explore the iron and myelin in DGM during aging. For iron deposition, the age-dependent paramagnetic susceptibility values in each DGM were fitted using an exponential model χ_*para*_ = *a*_1_ + *a*_2_*a*^−*a3y*^, where [*a*_1_, *a*_2_, *a*_3_] is the fitting coefficients array and *Y* is subjects’ ages. This model is similar to that used in the literature (Hallgren and Sourander, 1958). The Poisson fitting curve was used for fitting the diamagnetic susceptibilities in DGMs according to the previou study (Li et al., 2014) with a model of χ_*dia*_ = *a*_1_^y^2 + _3_, where [β_1_, β_2_, β_3_] is the corresponding fitting coefficients array.

### 3.5 Multiple-orientation imaging and sub-region segmentation

One representative healthy subject was scanned using a 3D GRE sequence at five different head orientations using a 3.0 T MR scanner (uMR790, UIH, Shanghai, China).

The 3D GRE scan parameters were: FOV = 240×240×160 mm^3^; matrix size = 240×240×80; spatial resolution = 1×1×2 mm^3^; TR = 35.0 ms; TEs = 4.5, 8.9, 13.4, 17.9, 22.4, 26.8 ms; FA = 15°; BW = 300 Hz/pixel; scan time = 7.7 min. The 2D MSE scan parameters were: FOV = 240×240 mm^2^; matrix size = 240×240; slice number = 80; spatial resolution: 1×1×2 mm^3^; TR = 4013 ms; TE_1_/spacing/TE_4_ = 16.72/16.72/66.88 ms; BW = 160 Hz/pixel; scan time = 17.8 min. The GRE magnitude images acquired at the tilted head orientation were linearly registered to GRE magnitude at the normal head position using FSL FLIRT (Jenkinson and Smith, 2001). The transformation matrix was then applied to the real and imaginary images. The five-orientation GRE images and corresponding *R_2_* map were combined in APART-QSM to produce high-quality QSM separation maps.

To demonstrate the superiority of the multiple-orientation QSM separation method, the multiple-orientation-based paramagnetic and diamagnetic susceptibility maps of the thalamus were merged into two parallel channels. The high-quality QSM maps were then compared with the manually segmented sub-regions of the thalamus, using the myelin-stained-section atlas as a reference (Schaltenbrand, 1977a). QSM, single-orientation APART-QSM, and multiple-orientation APART-QSM were compared for the definition of the tissue boundaries within the thalamus.

## 4. Results

### 4.1 Phantom validation

Fig. 1 illustrates the APART-QSM results of the agarose gel phantom with mixed susceptibility components. Fig. 1A presents the arrangement of phantom compositions as described in Section 3.2. Figs. 1B-1E show the susceptibility separation results of APART-QSM and χ-separation. χ_*para*_ maps (Figs. 1B, 1C) reconstructed from both APART-QSM and χ-separation successfully capture the regions where the cylinders are with only paramagnetic sources (the second row) and mixed susceptibility sources (the third row), but nearly zero susceptibility values in the cylinders with purely diamagnetic susceptibility sources in the first row. Similarly, χ_*dia*_ maps from the two methods exhibit comparable results (Figs. 1D, 1E). Specifically, the *ɑ*-map (Fig. 1F) obtained by APART-QSM exhibits varying values among different voxels. The linear regression analysis between susceptibility values from the single-source cylinders and those from mixed-susceptibility cylinders provide similar R-squared values in both methods (Figs. 1H, 1I). The slope of the linear regression from APART-QSM (0.989) is slightly higher than that from χ -separation (0.967). These results demonstrate that APART-QSM is able to accurately separate the opposing susceptibility sources in the mixed situation when the paramagnetic and diamagnetic susceptibility sources are ideally modeled as spheres in the agarose gel phantom. The reconstruction time of APART-QSM and χ-separation on the phantom data were 9.7 and 31.2 seconds, respectively.

### 4.2 In vivo brain susceptibility separation

Fig. 2 shows the reconstructed maps obtained by APART-QSM on one representative healthy subject. *M*_0_ is the extrapolated magnitude signal at TE = 0 ms based on the 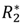 estimation of the first step in Eq. (5). The paramagnetic susceptibility source shows high positive values in χ_*para*_ map such as DGM, cortex, and blood vessels as well as some white matter regions. In contrast, χ_*dia*_ map reveals more negative susceptibility values in the white matter compared with DGM. Interestingly, the diamagnetic susceptibility values in the GP are also revealed by χ_*dia*_ map. Additionally, the magnitude decay kernels in the *ɑ*-map are close to the theoretical values of 323.5 Hz/ppm in most brain regions. However, decreased values of *ɑ*-map were found in the GP, the posterior limb of the internal capsule (PLIC), and the optic radiation, as pointed out by the red arrows.

**Figure 2.**
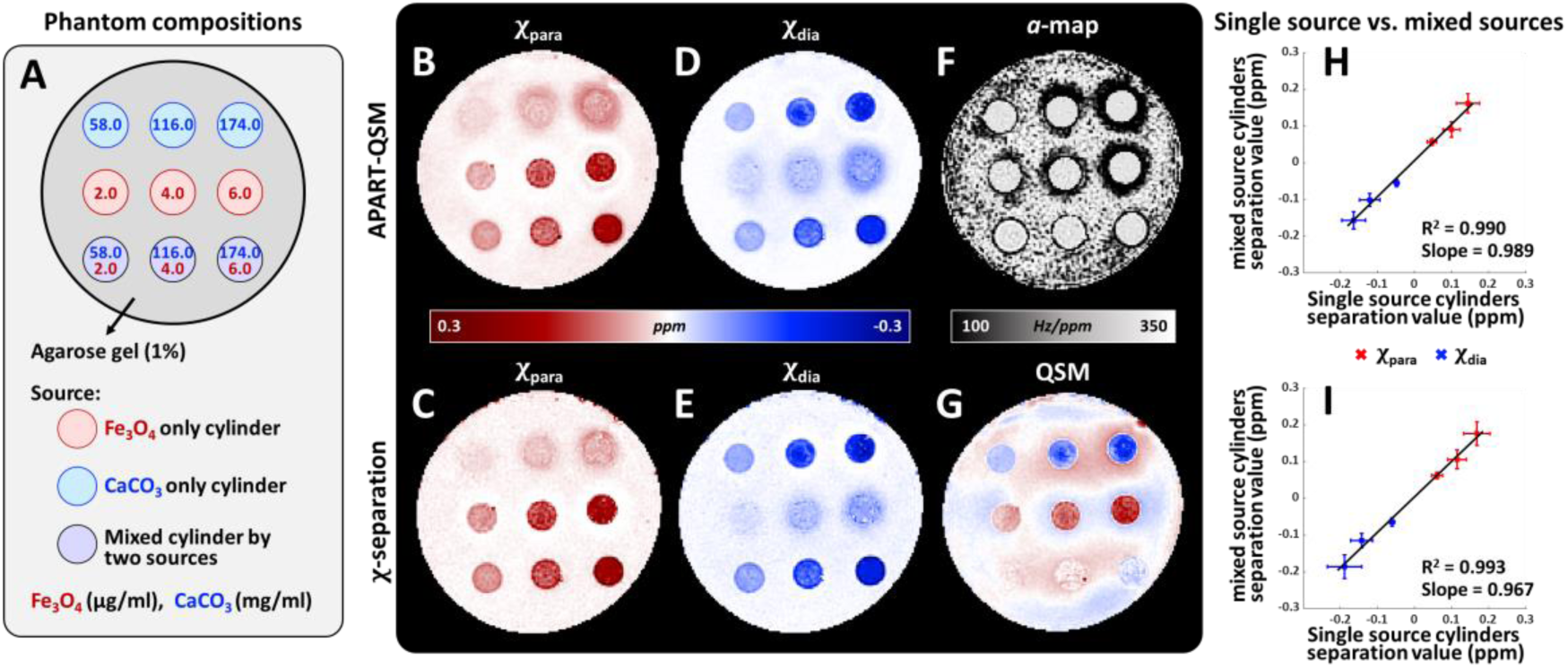
APART-QSM results of a representative healthy human subject. (A) The extrapolated signal *M*_0_ at TE = 0 ms. (B) The paramagnetic susceptibility map. (C) The diamagnetic susceptibility map. (D) The voxel-specific *ɑ*-map. (E) *ϕ*_*res*_denotes the time-independent residual phase. (F, G) 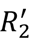 and 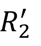 ɸ_*res*_ map representing the residual phase which is independent of echo time is shown in Fig 2E. 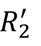 and 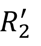 maps show clearer brain tissue boundaries than 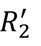 such as the outline of the thalamus indicated by the yellow arrows. The white matter fiber structures are also distinguishable from surrounding tissues on both χ_*dia*_ and 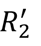 maps.

Fig. 3 shows the comparison of APART-QSM’s and χ-separation’s results on the healthy subject. Fig. 3A are two representative slices of QSM, χ_*para*_, and χ_*dia*_ maps reconstructed by the two methods. Visually, the χ_*para*_ map reconstructed by APART-QSM reveals a similar contrast to that obtained by χ-separation, whereas the χ_*dia*_ map from APART-QSM shows more negative susceptibility values in DGMs such as the GP and SN. Additionally, APART-QSM produces more organized white matter tissues in the χ_*dia*_ map than χ -separation. Figs. 3B-3D compare χ_*para*_, χ_*dia*_ maps with iron-, myelin-staining images in three brain regions outlined by the boxes in Fig. 3A. The χ_*para*_ map reconstructed from APART-QSM exhibits positive susceptibility values dominantly in the cortex and sparsely in the PLIC (yellow arrows), but nearly zeros in the prefrontal white matter (black arrows). While the χ_*para*_ map from χ-separation shows relatively higher positive susceptibility values in the prefrontal white matter area and lower susceptibility values in PLIC. Overall, the iron distribution in the prefrontal area revealed by χ_*para*_ map reconstructed from APART-QSM is more consistent with iron staining than that reconstructed from χ -separation. Similarly, the spatial variation of the diamagnetic susceptibility values in white matter revealed by APART-QSM’s χ_*dia*_ map matches well with the myelin-staining image. For example, the tissue boundaries between white matter (dark green arrows) and surrounding gray matter (light green arrows) in the prefrontal area are better delineated by APART-QSM than χ-separation. It is noted that the diamagnetic susceptibility values were clearly demonstrated in the GP, SN, and RN by APART-QSM’s χ_*dia*_ map, which is verified by the myelin staining as indicated by the orange, pink, and purple arrows in Figs. 3C & 3D. In comparison, the χ_*dia*_ map reconstructed by χ-separation shows barely diamagnetic susceptibility values in these DGMs. The reconstruction time of APART-QSM and χ-separation on this human brain data were 93.7 and 127.8 seconds, respectively.

**Figure 3.**
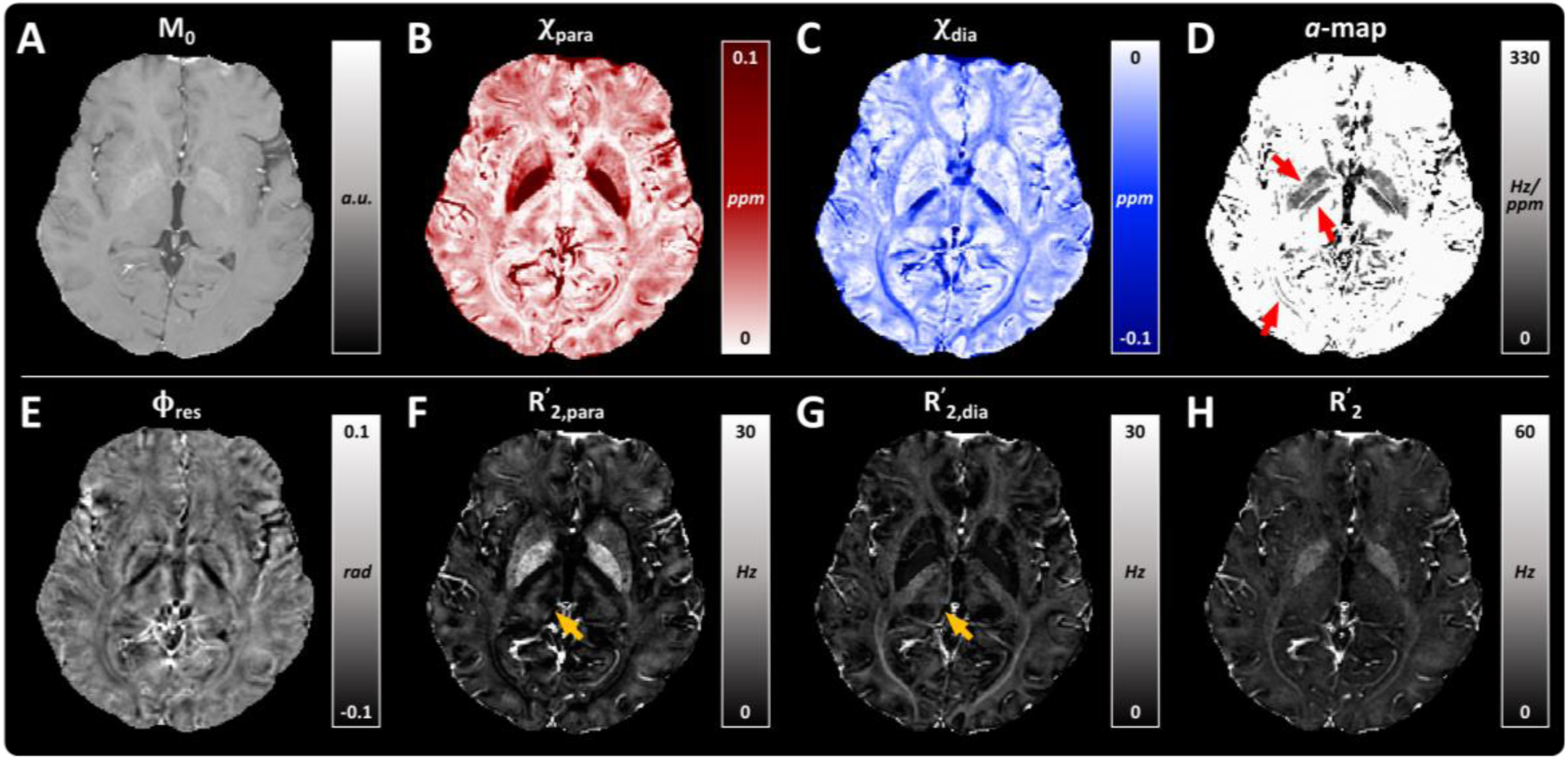
Comparisons of QSM, APART-QSM, and χ-separation on the representative healthy human brain data. (A) Two representative slices of QSM, paramagnetic and diamagnetic susceptibility maps from two methods. (B, C, D) Three brain regions outlined by the colored boxes in (A). The spatial variations of χ_*ɑrɑ*_ and χ maps reconstructed from APART-QSM are similar to the iron and myelin staining patterns. The reconstructed maps from APART-QSM show sparse positive values in the PLIC (yellow arrows) and clear diamagnetic susceptibility values in the GP (orange arrows), SN (pink arrow), and RN (purple arrow), but not by the reconstructed susceptibility maps by χ-separation. PLIC: posterior limb of the internal capsule; GP: globus pallidus; SN: substantia nigra; RN: red nucleus.

### 4.3 Quantitative analysis

Quantitatively, Fig. 4 presents the mean values of conventional QSM, χ_*para*_, χ_*dia*_ and composite QSM in five DGM nuclei averaged over 32 subjects. The reconstruction time of APART-QSM and χ-separation on these data were 86.3 ± 21.6 and 99.6 ± 3.2 seconds, respectively. Overall, APART-QSM and χ -separation both produce higher susceptibility values in these DGMs than conventional QSM. Additionally, χ-separation gives more paramagnetic susceptibilities in the GP, PUT, CN, and SN, while the proposed method yields relatively less paramagnetic susceptibilities in these regions. More negative susceptibility values are provided by APART-QSM than χ-separation in all five DGMs, especially in the GP and SN. The composite QSM by χ_*para*_ and χ_*dia*_ from APART-QSM shows comparable mean values to the conventional QSM, compared with that from χ-separation. These results may suggest that the proposed method can well preserve the intrinsic volume susceptibility values.

**Figure 4.**
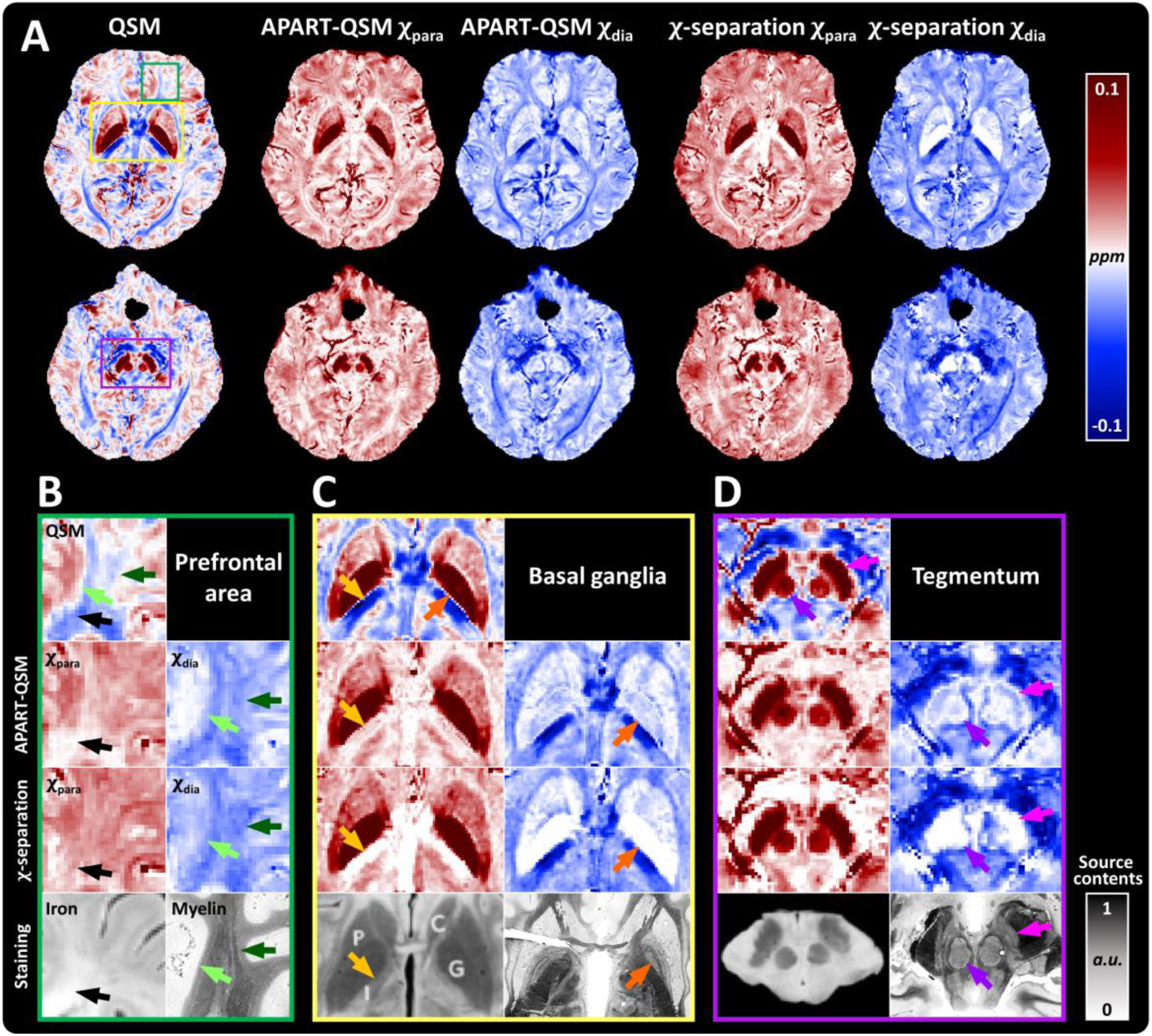
Region analysis in five selected DGMs of subjects (n=32) for QSM, paramagnetic and diamagnetic susceptibility maps, and composite QSM from APART-QSM and χ-separation. All data are presented as mean ± standard deviation. GP: globus pallidus; PUT: putamen; CN: caudate nucleus; SN: substantia nigra; RN: red nucleus; DGM: deep gray matter.

Fig.5 shows the quantitative analysis and comparison between APART-QSM and χ-separation of 32 subjects. Fig. 5A presents the ROIs of five DGM nuclei in myelin staining and QSM images. Fig. 5B reveals three age-different subjects’ QSM images in the MNI space after group-wise normalization for atlas construction. The DGMs in all subjects share consistent normalized locations, demonstrating the accuracy of constructed atlases. Fig. 5C shows the linear regression analysis between χ_*para*_ versus non-haemin iron concentrations in the GP, PUT, CN, SN, and RN for APART-QSM and χ-separation, respectively, where the iron concentrations are estimated from the fitted age-dependent curves in (Aoki et al., 1989; Hallgren and Sourander, 1958). The APART-QSM method achieves a slightly higher R-squared value than χ-separation. For the diamagnetic part shown in Fig. 5D, APART-QSM-based χ_*dia*_ atlas agrees well with the myelin staining images (https://brains.anatomy.msu.edu/) while χ-separation seems to underestimate the diamagnetic susceptibility in the GP and SN.

**Figure 5.**
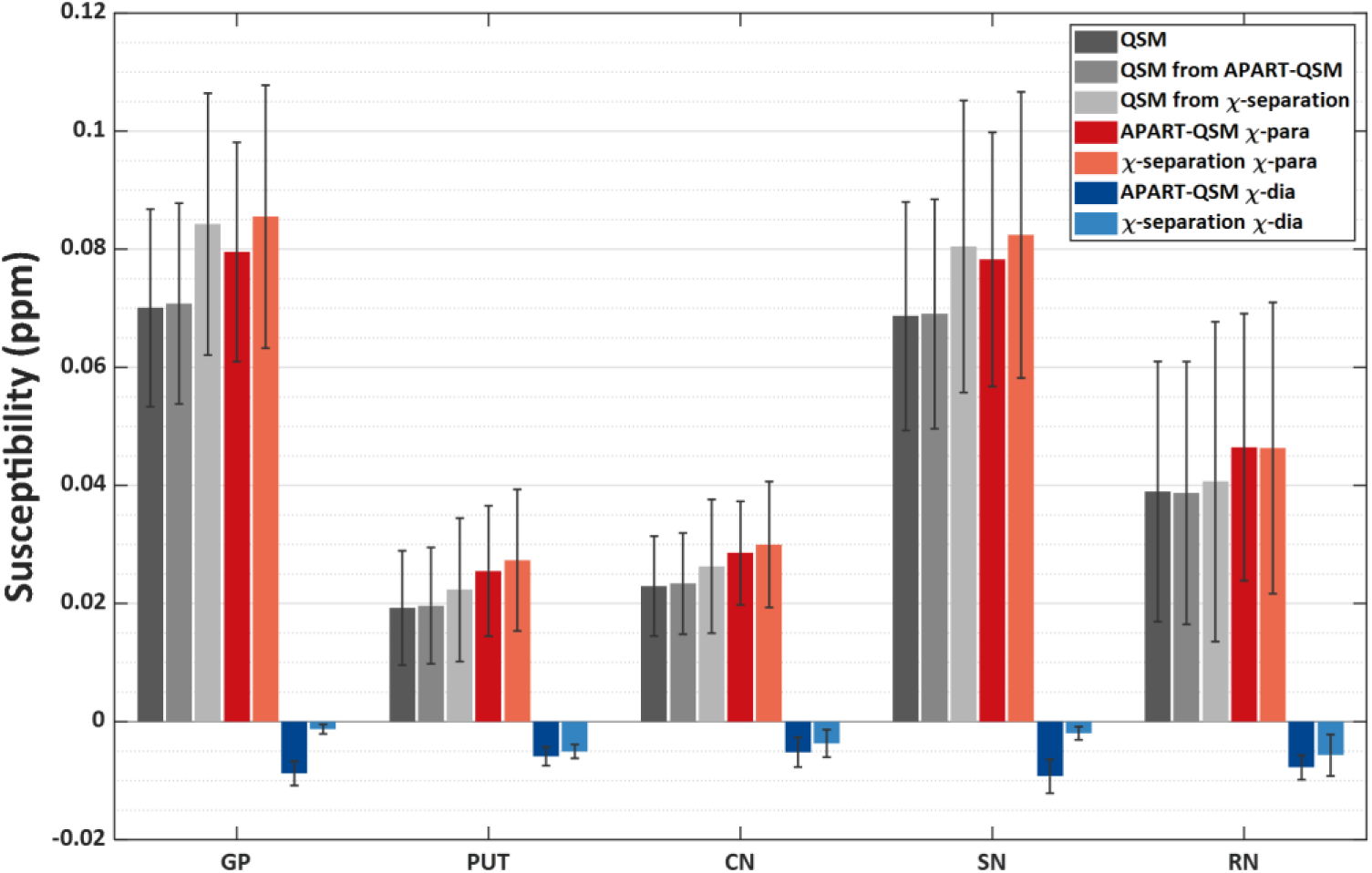
Quantitative validation on human subjects (n=32) and the subject-based atlas. (A) Five DGM labels drawn on myelin staining images and the built atlas. (B) Three age-different subjects’ QSM images in the MNI space. (C) Linear correlations between non-haemin iron concentration and χ_*ɑrɑ*_ values estimated by the two methods in DGMs. (D) Linear correlations between normalized myelin staining contents and χ_*ɑ*_ values from two methods in DGMs. GP: globus pallidus; PUT: putamen; CN: caudate nucleus; SN: substantia nigra; RN: red nucleus; DGM: deep gray matter.

### 4.4 Ex vivo brain imaging

Fig. 6 displays the comparisons of the reconstructed susceptibility maps, the *ɑ*-map from APART-QSM, and the susceptibility maps from χ -separation on one *ex vivo* macaque brain. The DTI’s fractional anisotropy (FA) map of the corresponding slice is also displayed. The stronger anisotropy implies more nerve fibers, with a simple linear relationship with more diamagnetic myelin in general. The χ_*dia*_ map from APART-QSM (Fig. 6B) shows a similar tissue contrast with the FA map, as pointed out by the arrows. While this continuity of FA can hardly be observed in the white matter region on the χ_*dia*_ map reconstructed from χ-separation (Fig. 6E). These results indicate that APART-QSM can successfully recover the distribution of diamagnetic susceptibility source in the brain white matter. <u>The reconstruction time of APART-QSM and χ-separation on this *ex vivo*</u> <u>data were 8625.9 and 8450.2 seconds, respectively.</u>

**Figure 6.**
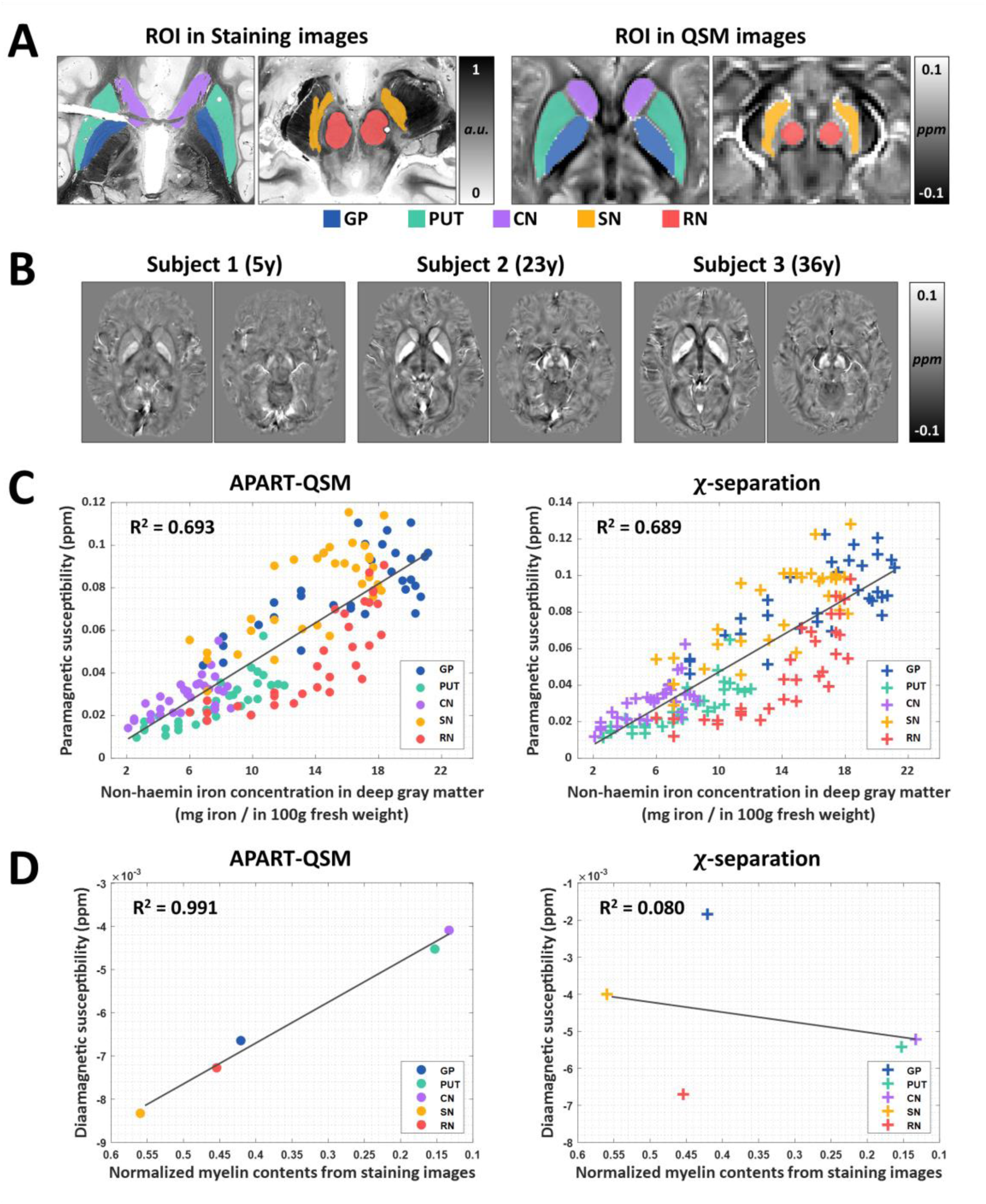
Qualitative comparison of the results of APART-QSM and χ-separation on one *ex vivo* macaque brain. (A-C) Paramagnetic and diamagnetic susceptibility maps reconstructed by APART-QSM, as well as the *ɑ*-map. (D, E) Two susceptibility maps reconstructed by χ-separation (F) The FA map from DTI. FA: fractional anisotropy.

Fig. 7 shows a more comprehensive comparison between the reconstructed diamagnetic susceptibility values with the myelin staining content. Eight labels in the white matter fibers were drawn on the normalized myelin staining images and diamagnetic susceptibility maps according to a parcellation of the macaque brain atlas (Fig. 7C) (Feng et al., 2017). The correlations between the mean diamagnetic susceptibility versus normalized myelin staining content within each ROI were illustrated in Fig. 7G & 7H. A higher R-squared value of 0.854 between staining content and χ_*dia*_ map from the APART-QSM was found than χ -separation (0.530), indicating the diamagnetic susceptibility estimated by APART-QSM is more consistent with myelin staining results.

**Figure 7.**
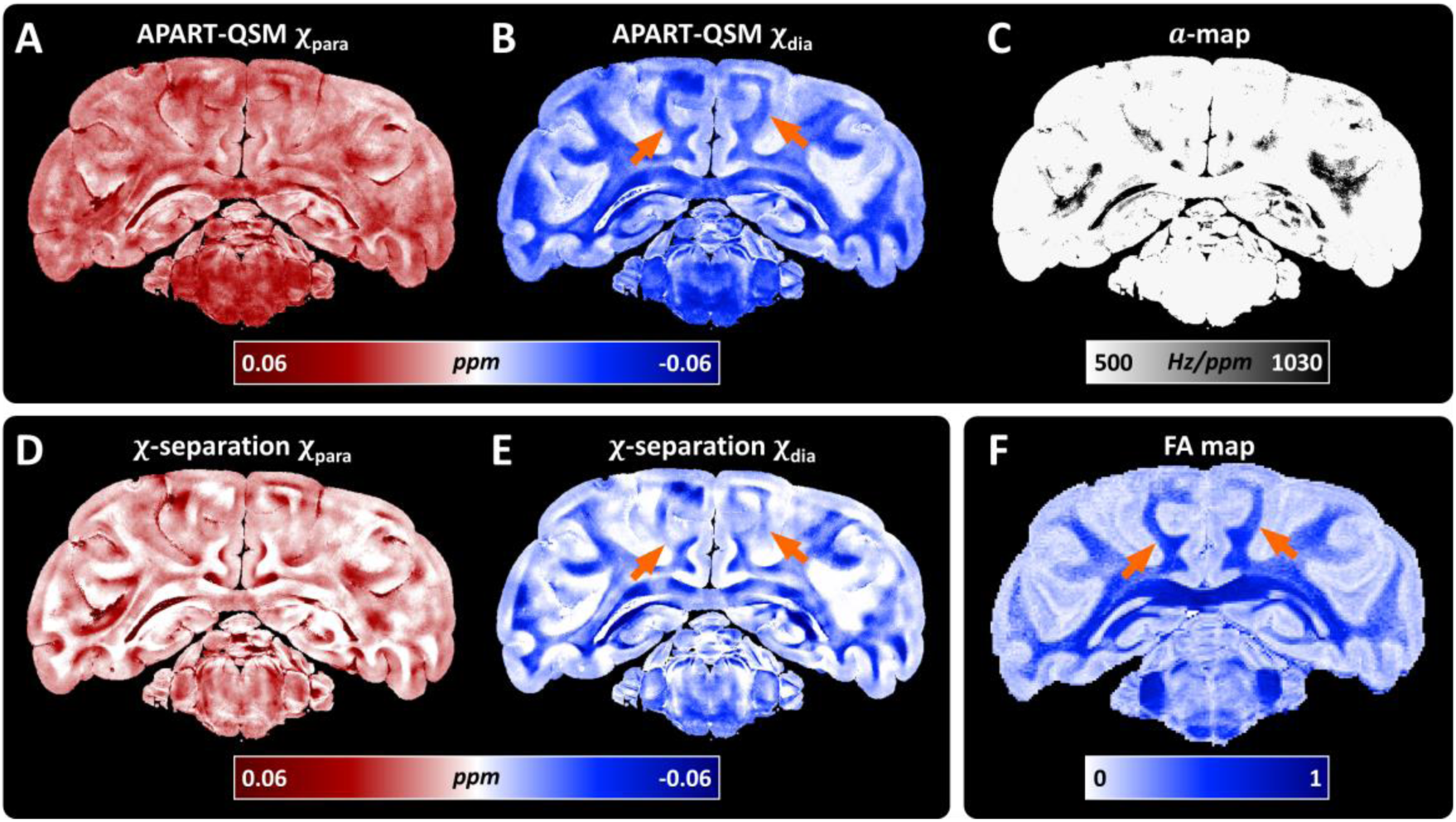
Quantitative comparisons on *ex vivo* macaque data. (A, B) Myelin staining image and corresponding normalized gray-scale map with labels. (C) A macaque brain atlas with eight white matter labels from literature. (D) Manual segmented ROIs on QSM referring to the macaque brain atlas. (E, F) Diamagnetic susceptibility maps from APART-QSM and χ-separation, respectively. (G, H) The linear regression analysis between ROI mean values of diamagnetic susceptibility from the two methods and normalized myelin staining content.

### 4.5 Longitudinal brain development and sub-region segmentation

The separation of paramagnetic and diamagnetic susceptibilities by APART-QSM provides the potential to quantify the brain iron and myelin changes simultaneously during aging. The age-dependent susceptibility maps (32 subjects) were separated by APART-QSM. Fig. 8 shows the susceptibility changes in the GP, SN, and PUT as a function of age using QSM, χ_*para*_ and χ_*dia*_ maps of all subjects. Overall, the QSM and χ_*para*_ maps’ susceptibility values in the GP and SN grow rapidly from 4 to 20-year-old then reach the plateaus around 30-year-old. The two maps’ susceptibility values in the PUT show continuous growth with the year of age until 39 years old. However, the fitted curves using conventional QSM exhibit underestimated susceptibility values than the reconstructed paramagnetic susceptibilities. In the GP, the mean diamagnetic susceptibility value gradually decreases (more diamagnetic) from 4 to 15 years, then reaches its minimum at around 20 years of age. In the SN, diamagnetic susceptibility decreases more rapidly from 4 to 20 years and reaches its peak diamagnetic values around 25 years of age. Differently, the diamagnetic susceptibility in the PUT maintains the trend of decreasing until 39 years old. These results demonstrate that the diamagnetic susceptibility components provided by APART-QSM have the potential to monitor the specific myelination and demyelination in different DGMs occurred during brain development and aging.

**Figure 8.**
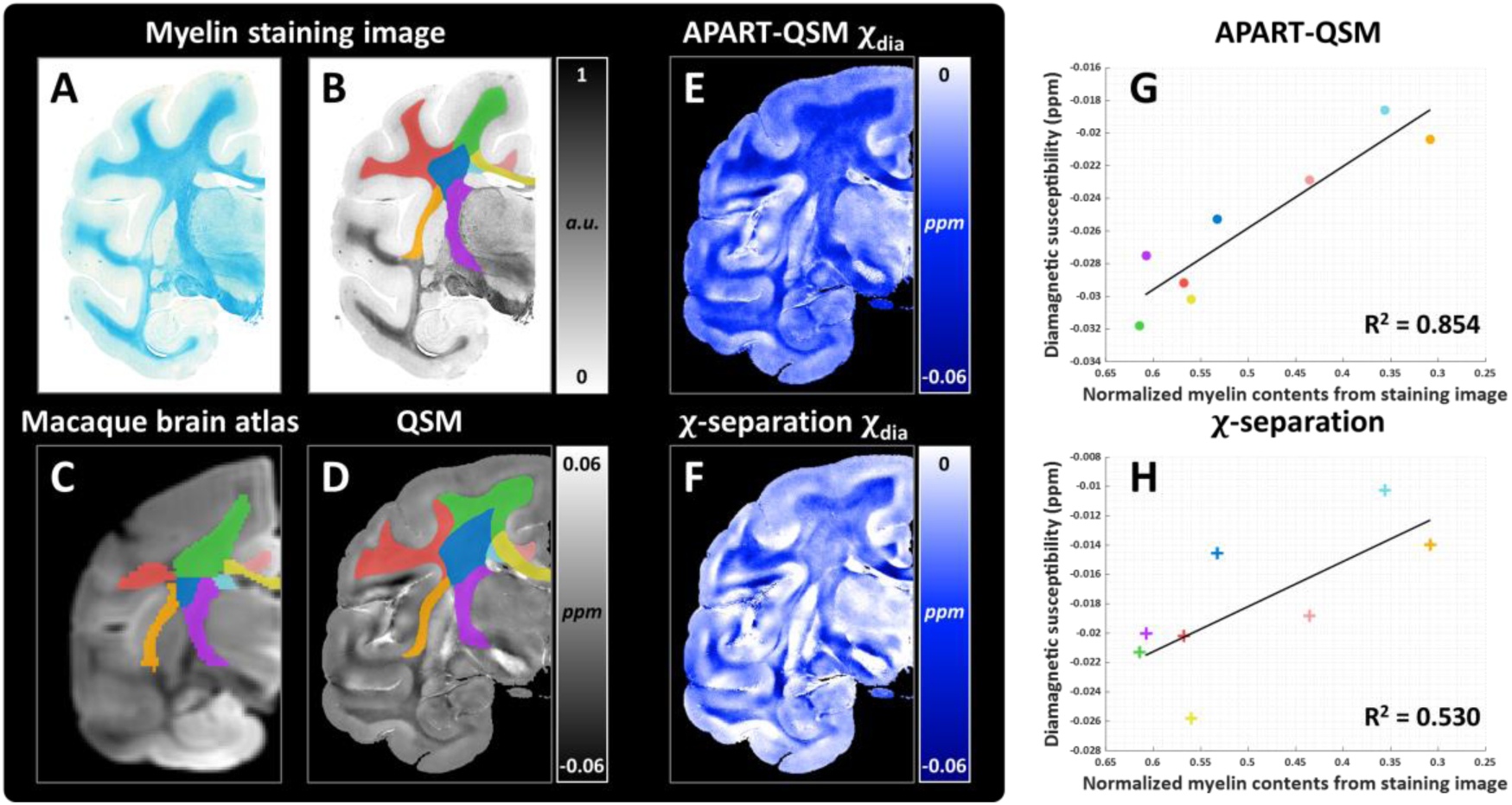
Susceptibility changes during brain development in the GP, SN, and PUT were investigated using QSM, χ_para*ɑrɑ*_ and χ_para*ɑ*_ maps reconstructed from APART-QSM. The solid and dashed curves represent the fitted developing trajectory and 95% prediction intervals, respectively. GP: globus pallidus; SN: substantia nigra; PUT: putamen.

We demonstrate the proposed method with multiple-orientation GRE data as input to provide high-quality susceptibility separation results. Fig. 9 shows the comparison of sub-region susceptibility contrast between conventional QSM (Fig. 9A), the results of APART-QSM using single-orientation (Fig. 9B) and multiple-orientation data (Fig. 9C).

**Figure 9.**
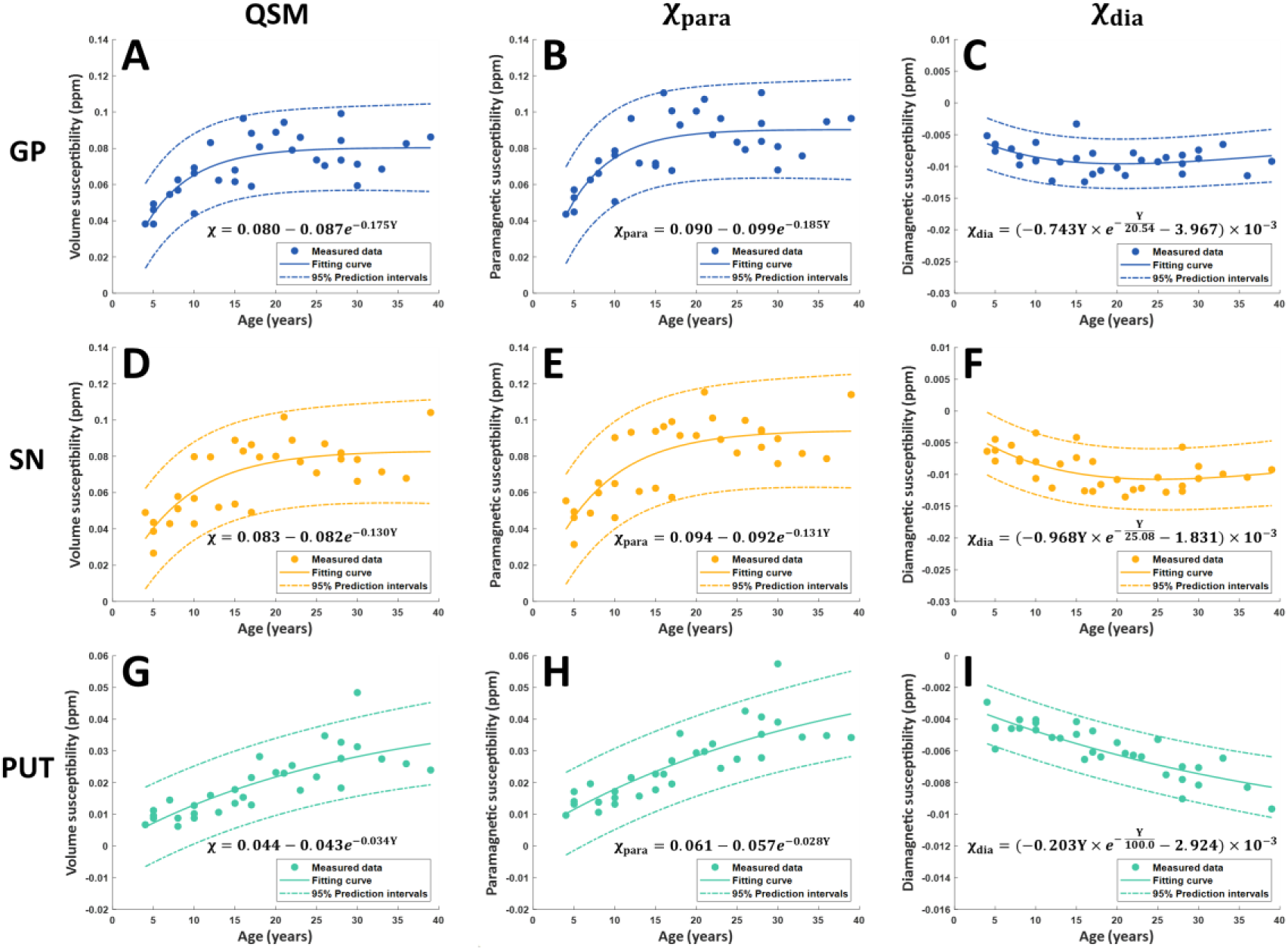
Sub-region segmentation of thalamus using merged χ_*paraɑrɑ*_ and χ_*paraɑ*_ maps reconstructed by multiple-orientation-data-based APART-QSM compared with QSM and single-orientation-data-based APART-QSM results. The merged χ_dia*ɑrɑ*_ and χ_*diaɑ*_ maps are set in red and green channels, respectively. (A) Conventional QSM. (B) Single-orientation-based APART-QSM results (C) Multiple-orientation-based APART-QSM results and fine segmentation in the thalamus (see the Supporting materials for details of sub-nuclei in the thalamus). (D) Myelin staining images and fine segmentation in the thalamus. PUT: putamen; GP: globus pallidus.

The χ_*para*_ and χ_*dia*_ maps were merged into parallel channels for better visualization (see the two-dimensional color bar in Fig. 9). The QSM map only shows the overall outline of the thalamus; the separation results using single-orientation data appear to have more diamagnetic susceptibility values, as pointed out by yellow arrows, whereas there are no clear boundaries of sub-region in the thalamus. In contrast, the separation results using multiple-orientation data show great diamagnetic susceptibility details and tissue edges, especially in medial thalami (M) and pulvinar thalami (Pu) as pointed out by white arrows and reach feasibility for sub-region segmentation, matching with the manually segmented sub-regions defined based on the staining intensity (Fig. 9D). The reconstruction time of APART-QSM on this multiple-orientation data was 369.9 seconds.

## 5. Discussion

QSM is a promising non-invasive MR technique that quantifies tissue volume magnetic susceptibility. However, conventional QSM methods only provide the voxel-averaged susceptibility value with mixed information from opposing susceptibility sources in a single voxel, restricting the further potential applications of QSM. This study proposed a new magnetic susceptibility separation method called APART-QSM, which utilizes a comprehensive complex model with an iterative data fitting algorithm to accurately separate intravoxel paramagnetic and diamagnetic susceptibility. Results from phantom, *ex vivo* macaque brain, and *in vivo* human brain experiments demonstrate that the proposed method provides superior performance for separating the opposing magnetic susceptibility sources than the compared methods. In addition, the proposed APART-QSM was applied to the investigation of longitudinal brain development and deployed with multiple-orientation data to provide high-quality isolated susceptibility maps for a better description of brain nuclei sub-regions. These results indicate that the proposed APART-QSM would broaden our understanding of tissue magnetic susceptibility properties in the field of neuroscience.

### 5.1 Motivation for proposing the spatially varying magnitude decay kernel

The proposed APART-QSM improves the QSM separation accuracy, which benefits from the more comprehensive complex signal model and the spatially variable magnitude decay kernel solved by the proposed alternate fitting scheme. Previous studies model the susceptibility sources as uniformly magnetized spheres whose MRI signals evolve under the static dephasing regime (Chen et al., 2021; Emmerich et al., 2021; Shin et al., 2021). This assumption results in a uniform magnitude decay kernel (parameter *ɑ*) over the whole brain tissue. The spatial-invariant parameter *ɑ* was generally pre-estimated by the slope of the linear regression between 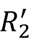 relaxation and susceptibility values only in a few DGMs based on the assumption that these regions are primarily dominated by paramagnetic iron. However, the calculation and assumption about the magnitude decay kernel are too ideal for conforming to the real biochemical environment. First, the myelin staining images in Fig. 3 indicate that DGMs such as the GP, SN, and RN all contain non-neglectable diamagnetic sources. Thus, the parameter *ɑ* pre-determined in these regions might be inaccurate. Second, the static dephasing regime ignores the geometry of the microstructure, the size and concentrations of sources (Brammerloh et al., 2021; Duyn and Schenck, 2017; Yablonskiy et al., 2021), and the water diffusion effect where the molecule interchange during diffusion affects the phase accumulation and causes irreversible MRI signal loss in *in vivo* biological tissues (Yablonskiy and Haacke, 1994). These facts indicate that a fixed spatial-invariant magnitude decay kernel is insufficient to characterize the relationship between 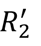 relaxation and susceptibilities due to the complex microenvironment in biological tissues. To this end, APART-QSM regards the magnitude decay kernel (i.e., parameter *ɑ*) as voxel-specific, however, it can more realistically characterize the 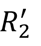 relaxation behavior under the susceptibility effect. Furthermore, the proposed iterative fitting algorithm enables updating between the parameter *ɑ* and susceptibilities, preventing the errors of the independently estimated parameter *ɑ* from propagating to the final QSM separation reconstruction.

### 5.2 Selection of regularization parameters and the initialization

The APART-QSM method relies on the high-quality estimation on 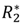 and 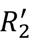. The choice of echo time mainly depends on the tissues’ susceptibility and *T*^∗^ relaxation properties. In the phantom experiment, faster signal attenuation occurs due to the larger magnetic field inhomogeneity caused by large susceptibility sources. Therefore, the maximum echo time of 14.7 ms was selected to avoid signal attenuation under the noise level, while ensuring that the 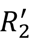 ( = 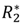 − *R_2_*) can be accurately calculated from measured 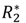 and *R_2_* signals. On the other hand, the bio-tissues appear to have relatively lower susceptibility effects than phantom fabricated with high concentrations of Fe_3_O_4_ or CaCO_3_. Thus, longer TEs are used for brain imaging to obtain more phase accumulation, consequently improving the QSM reconstruction accuracy.

The initialized pre-calculated QSM map was reconstructed by STAR-QSM as a constraint on the susceptibility range for the sum of the opposing susceptibility components. As shown in Fig. S1, the χ_*para*_ and χ_*dia*_ maps of the phantom obtained from the STAR-QSM as the initialization show the most realistic contrast with high susceptibility quantification accuracy. The over smooth effect caused by the highly regularized QSM reconstruction could decrease the reconstruction performance. Other methods, such as the FANSI method with magnitude-weighted non-linear total variation (nlTV_mag) (Milovic et al., 2018; Papadakis et al., 2015), could also be used as the initialization, and similar results were obtained as demonstrated in Fig. S2. In the future, more efforts will be needed to develop more advanced QSM separation methods without the pre-determined QSM as the susceptibility range constraint.

The first two regularization parameters (λ_1_ and λ_2_) were determined based on the L-curve method tested on a representative healthy human brain data (Fig. S3). λ_1_ = 0.1 and λ_2_ = 10 were found at the corner of the corresponding L-curves. TV regularization was used for noise suppression. The L-curve indicates that *λ*_3_ = 8 seems to be a corner point (Fig. S4). However, the reconstructed χ_*para*_ and χ_*dia*_ maps become over-smooth when *ʎ*_3_ = 8, which might be due to the low-level noise in the *in vivo* data. On the other hand, the TV regularization is to suppress the noise when facing high-resolution data with a lower signal-to-noise ratio. Therefore, *λ*_3_ = 1 was selected as an appropriate parameter of the TV regularization to balance noise suppression and quantification accuracy. In addition, TV regularization can be regarded as an independent condition in the proposed model. This could avoid the under-fitting situation because of the limited echoes. Other constraints, such as nonlinear TV and sparsity, can be also integrated into the model.

### 5.3 Results analysis

The magnitude decay kernel (*ɑ*-map) from our method in the phantom experiment reveals a small spatial variation (Fig. 1F). This is likely because the phantom experiment only simulates a relatively simple situation of sources’ mixture with ideal spherical susceptibility sources (Fe_3_O_4_, CaCO_3_) and nearly isotropic diffusion effects. Despite limited variations, the fine-tuned *ɑ*-map enables APART-QSM to obtain more reasonable χ_*para*_ and χ_*dia*_ maps, achieving a high linear regression result whose slope is closer to 1 than that from χ-separation method (Fig. 1H & 1I). Low *ɑ* values appear surrounding the cylinder with a single high-concentration source. This is likely coming from imperfect dipole inversion of the susceptibility with a sharp transition between the tube boundary and the background of the phantom. However, this inaccuracy is local and does not affect the estimation of *ɑ* value and susceptibility in the cylindrical regions. The artifacts could not happen for the *in vivo* and *ex vivo* brain data since bio-tissues should not have sharp edges.

Due to the spatial-variant *ɑ*-map offering a more realistic relationship between 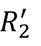 relaxation and absolute susceptibility, the qualitative and quantitative comparisons (Fig. 3 & 5) with histological studies (Drayer et al., 1986; Naidich et al., 2013) demonstrate successful separation of paramagnetic and diamagnetic components by APART-QSM. More importantly, the fine-turned *ɑ*-map enables the subtle myelin fibers in DGMs to be distinguishable from dominant iron in DGMs (Haider et al., 2014) (Fig. 8C, 8F, 8I), providing the potential for quantifying the myelination in brain DGMs. We thus believe that our proposed method greatly improves the accuracy of quantifying the paramagnetic and diamagnetic sources of brain tissue.

The susceptibilities reconstructed from the single orientation phase data were still contaminated by streaking artifacts due to the ill-posed nature of QSM (Deistung et al., 2017). To resolve this issue, multiple-orientation phase data were deployed in APART-QSM to stabilize the inversion relating the tissue phase to the underlying magnetic susceptibility values. The merged map of χ_*para*_ and χ_*dia*_ reconstructed by multiple-orientation APART-QSM better resembles the myelin-staining-based atlas (Schaltenbrand, 1977b) than single-orientation APART-QSM results, as illustrated in Fig. 9. These results demonstrate that multiple-orientation APART-QSM can significantly improve performance for the delineation and segmentation of small-size brain structures (e.g., sub-structures of the thalamus).

### 5.4 Potential applications

The proposed method non-invasively quantifies the intravoxel paramagnetic and diamagnetic susceptibilities of the *in vivo* human brain. It may help us to understand the sophisticated relationship between brain iron and myelin during normal brain aging and in various brain disorders. For instance, iron deposition and myelination/demyelination are both associated with the normal aging process (Fig. 8) and neurodegenerative diseases (Bartzokis, 2004; Ward et al., 2014). A previous study has indicated that the myelinated fibers in DGMs can be visualized and quantified by a susceptibility tensor imaging (STI) technique based on acquired multi-orientation data (Bao et al., 2021). APART-QSM could be a novel imaging technique for quantifying the subtle magnetic susceptibility of myelin in brain DGMs based on QSM. It could open a new window to monitor the myelination and demyelination process in brain DGMs. Furthermore, accurate mapping of the iron and the myelin distribution in the cortical and subcortical structures may facilitate accurate *in vivo* atlases of iron and myelin across the lifespan. The built atlas can be used to indicate abnormal alterations associated with related brain disorders, such as Alzheimer’s disease (Bulk et al., 2018; Liu et al., 2018), Parkinson’s disease (Dean et al., 2016; Foley et al., 2022), and multiple sclerosis (Hametner et al., 2013). Additionally, the multiple-orientation APART-QSM achieves a clear boundary definition of small brain nuclei, which may help improve the accuracy of tissue segmentation and visualization. Clinically, direct target localization of brain nuclei is extremely helpful for deep brain stimulation (DBS) planning. For example, the Vim in the thalamus, which can be observed in the results of multiple-orientation APART-QSM, is a potential target for the treatment of essential tremors (Papavassiliou et al., 2004).

### 5.5 Limitations

There are some limitations in our work. First, simulating the complicated sources mixture in the phantom experiments is challenging. Only simple mixed susceptibility sources were performed. In addition, an extra T2 mapping scan is still employed in our model to estimate the *R_2_* map, which will increase the total acquisition time of MR scanning and potential error from inter-scan head motion. Recent proposed multi-contrast imaging techniques, such as echo planar time-resolved imaging (Wang et al., 2019) or multi-tasking MRI (Cao et al., 2022) with simultaneously T2* and T2 imaging, could be alternatives to overcome these problems.

## 6. Conclusion

The separation of paramagnetic and diamagnetic susceptibility would be of utmost importance for broadening QSM applications. We proposed the APART-QSM method for improved susceptibility sources’ separation using comprehensive complex data modeling and iterative voxel-specific magnitude decay kernel estimating algorithm. The calculated voxel-wise magnitude decay kernel could realistically model the relationship between the 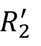 relaxation and absolute susceptibility. Phantom and *ex vivo* experiments demonstrate excellent performance for simultaneously quantifying brain iron and myelin. The results on *in vivo* human data further illustrate the superiority of our method in subtle source extraction, such as imaging the myelin fibers in DGM and iron in white matter. Furthermore, the proposed multiple-orientation APART-QSM provides a more faithful tissue delineation of the small brain regions. A longitudinal study using the proposed method may bring novel insights into the internal neural mechanisms for brain development and aging. We thus believe that APART-QSM could be helpful in quantifying the abnormal alterations of tissue magnetic susceptibility associated with various brain disorders in the future.

## Acknowledgments

This study is supported by the National Natural Science Foundation of China (61901256, 91949120, 62071299) and by the Shanghai Science and Technology Development Funds (21DZ1100300).

## Appendix

### The complex-valued GRE signal model

The raw complex GRE signal *S* in position *r* can be written:

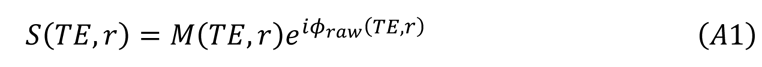

where *M* is the magnitude image, affected by the 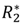 (= *R_2_* + 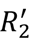) relaxation. *ϕ*_*rɑ*_ is an echo time-dependent phase signal and *TE* is the echo time. The magnitude and phase can be expressed as:

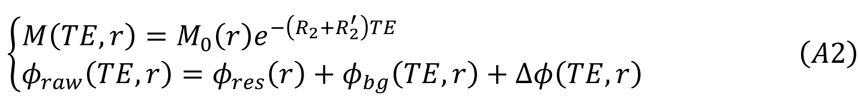

where *M*_0_ is the extrapolated magnitude signal at TE = 0 ms. *R_2_* is the natural relaxation rate and 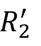 denotes the extra relaxation rate attributed to the field inhomogeneity. ɸ_*res*_ represents the time-independent residual phase. *ϕ*_*TE*_ is the background phase and Δ*ϕ* denotes the phase shift caused by susceptibility sources. Therefore, the complex signal *S* can be expressed as:

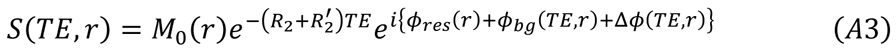

When ignoring the diffusivity and sources’ anisotropy in the theory of static dephasing (Yablonskiy and Haacke, 1994), the transverse relaxation rate 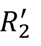 linearly relates to the absolute susceptibility effects from opposite sources. In addition, the phase shift Δ*ϕ* can be expressed as the signed sum of the opposite susceptibility sources. Thus, the susceptibility separation model at *j*^th^ echo in the static magnetic field *B*_0_ can be further expanded as:

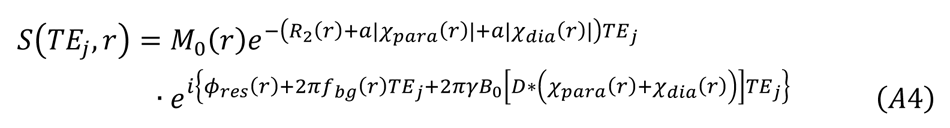

where *ɑ* is the magnitude decay kernel. χ_*para*_ and χ_*dia*_ are the opposite susceptibility sources. 2Π*f*_*bg*_*TE*_*j*_ denotes the background phase. *γ* is the gyromagnetic ratio and *D* is the magnetic dipole kernel.

