## Supplemental materials for "APART-QSM: an improved sub-voxel quantitative susceptibility mapping for susceptibility source separation using an iterative data fitting method"

**Supporting materials**

**Labels in the thalamus in Figure 9:**

A.m: nucleus anteromedialis thalami;

B.co.s: brachium colliculi superioris;

Ce: nuclei centrales thalami;

Ce.mc: nucleus centralis magnocellularis;

Cm.hb: commissura habenularis;

Fa: nucleus fasciculosus thalami;

La.m.ip: lamella medialis interpolaris;

La.m.o: lamella medialis oralis;

Li: nucleus limitans thalami;

Lpo: nucleus lateropolaris thalami;

M: territorium mediale thalami;

Pf: nucleus parafascicularis thalami;

Pu: pulvinar thalami;

S.pv: substantia periventricularis;

T.mth: Tractus mamillo-thalamicus;

V.c: nuclei ventrocaudales;

V.im: nuclei ventrointermedii;

V.o: nuclei ventroorales;

Z: territorium zentrolaterale thalami.

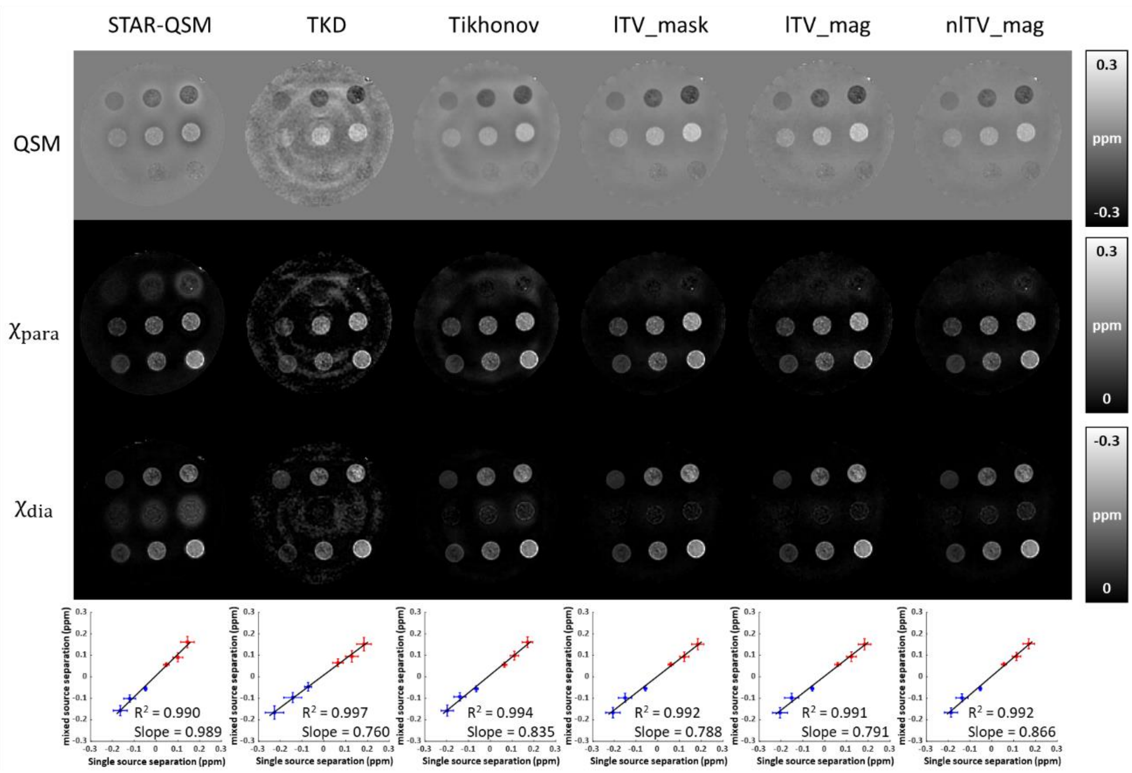


**Figure S1.** Susceptibility separation results from six different initialization schemes on the phantom data. TKD: thresholded k-space division; Tikhonov: closed-form reconstruction for L2-regularized QSM; lTV_mask: brain mask-weighted linear total variation; lTV_mag: magnitude-weighted linear total variation; nlTV_mag: magnitude-weighted non-linear total variation.

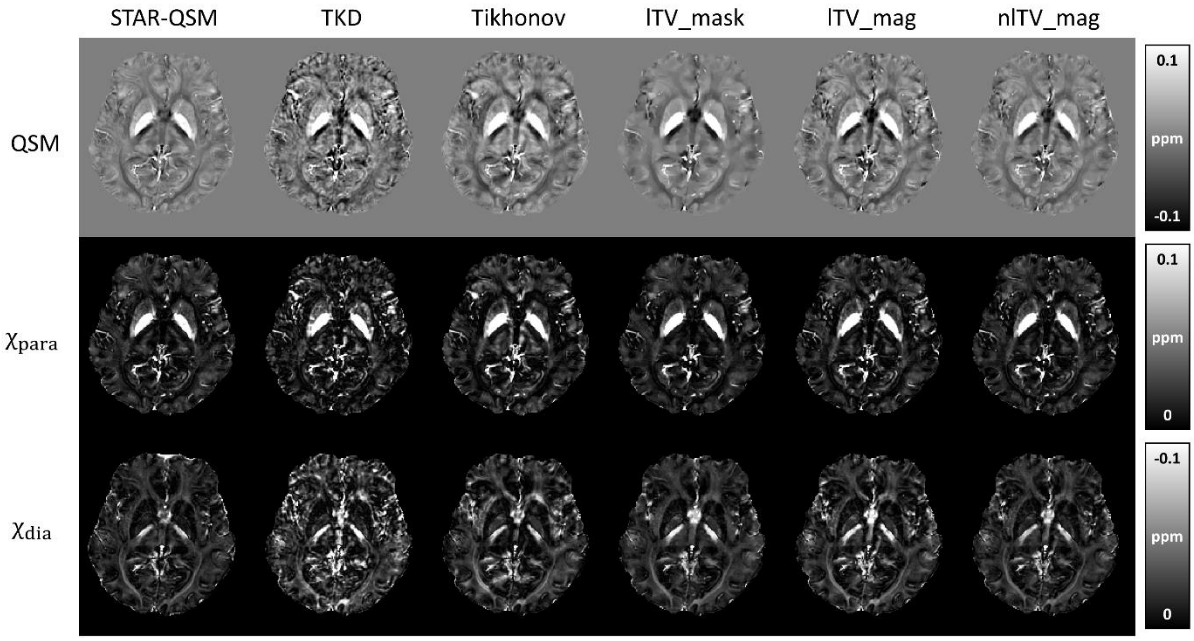


**Figure S2.** Susceptibility separation results from six different initialization schemes on the *in vivo* data. TKD: thresholded k-space division; Tikhonov: closed-form reconstruction for L2-regularized QSM; lTV_mask: brain mask-weighted linear total variation; lTV_mag: magnitude-weighted linear total variation; nlTV_mag: magnitude-weighted non-linear total variation.

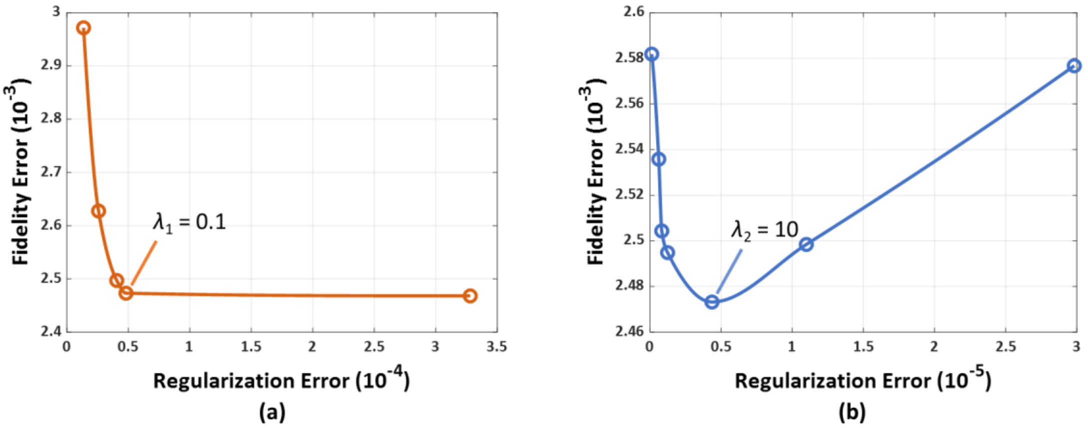


**Figure S3.** Selection of the optimal regularization weightings, $\lambda_1$ and $\lambda_2$, according to the L-curves. (a) The fidelity error versus the regularization error when $\lambda_1$ varying from 0.05 to 0.4 with $\lambda_2$ and $\lambda_3$ fixed to be 10 and 1. (b) The fidelity error versus the regularization error when $\lambda_2$ varying from 1 to 30 with $\lambda_1$ and $\lambda_3$ fixed to be 0.1 and 1.

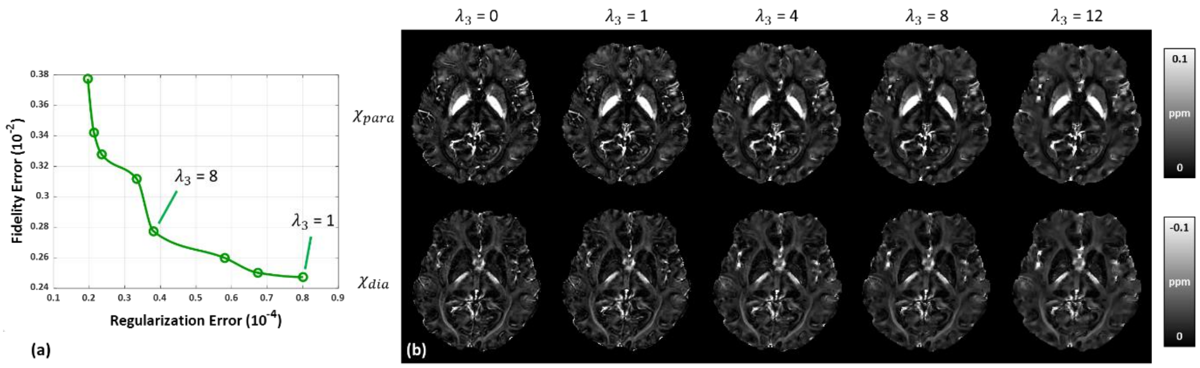


**Figure S4.** Selection of the optimal total variation regularization parameter $\lambda_3$. (a) The fidelity error versus the regularization error when $\lambda_3$ varying from 1 to 20 with $\lambda_1$ and $\lambda_2$ fixed to be 0.1 and 10. (b) Reconstructed $\chi_{para}$ and $\chi_{dia}$ maps under different $\lambda_3$ values.

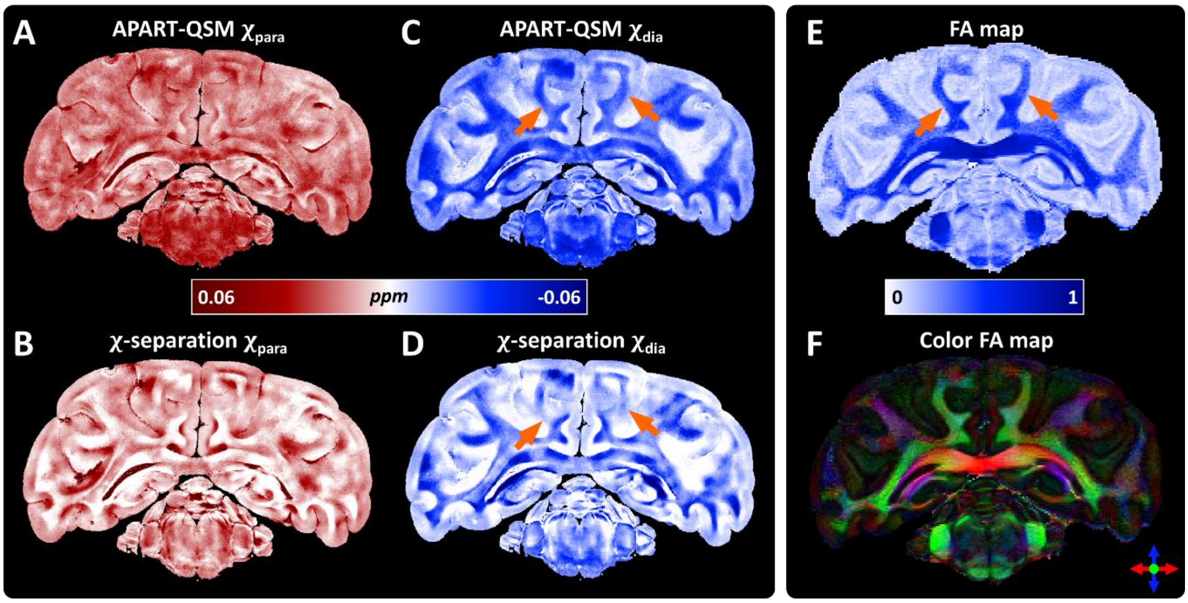


**Figure S5.** The results of APART-QSM and $\chi$-separation compare with FA maps from DTI on one *ex vivo* macaque brain. (A-D) Paramagnetic and diamagnetic susceptibility maps reconstructed by APART-QSM and $\chi$-separation (E) The FA maps. (E) The directional (color) FA maps. FA: fractional anisotropy.
